# Adapting CRISPR/Cas9 System for Targeting Mitochondrial Genome

**DOI:** 10.1101/2020.02.11.944819

**Authors:** Syed-Rehan A. Hussain, Mehmet E. Yalvac, Bendict Khoo, Sigrid Eckardt, K. John McLaughlin

**Author notes:** Corresponding Authors S-R A Hussain, Center for Clinical and Translational Research, Abigail Wexner Research Institute at Nationwide Children’s Hospital, 700 Children’s Drive, Columbus, OH 43205, USA., K. J McLaughlin, Center for Molecular and Human Genetics, Abigail Wexner Research Institute at Nationwide Children’s Hospital, 700 Children’s Drive, Columbus, OH 43205, USA. BK: Environmental Health Program, School of Public Health, University of Minnesota, Minneapolis, Minnesota, USA.

## Abstract

Gene editing of the mitochondrial genome using CRISPR-Cas9 system is highly challenging mainly due to sub-efficient delivery of guide RNA and Cas9 enzyme complexes into mitochondria. In this study, we were able to perform gene editing in the mitochondrial DNA by appending NADH-ubiquinone oxidoreductase chain 4 (ND4) targeting guide RNA to a RNA transport derived stem loop element (RP-loop) and expressing the Cas9 enzyme with preceding mitochondrial localization sequence. Our results showed mitochondrial co-localization of RP-loop gRNA and a marked reduction of ND4 expression in the cells carrying a A11204G variant in their ND4 sequence coincidently decreasing the mtDNA levels. This proof-of-concept study suggests that stem loop element added sgRNA can be transported to the mitochondria and functionally interact with Cas9 to mediate sequence specific mtDNA cleavage. Using this novel approach to target the mtDNA, our results provide further evidence that CRISPR-Cas9 mediated gene editing might potentially be used to treat mtDNA related diseases.

## INTRODUCTION

Mitochondrial diseases caused by defects in mitochondrial oxidative phosphorylation (OXPHOS) can be linked to the presence of wild type and mutant variants of mitochondrial genes in the cells, which is called heteroplasmy. Dependent on levels of heteroplasmy, they can manifest as highly heterogeneous disorders resulting in phenotypes ranging from mild hearing impairment to severe progressive multisystem disorders (1, 2, 3). Genetically, mitochondrial disorders typically include those resulting from variants in the mitochondrial genome (mtDNA) or those associated with alterations in nuclear genes coding for mitochondrial proteins. As many as 1:200 individuals harbor a pathogenic mtDNA variant that could be transmitted to offspring (4), and the population prevalence for specific pathogenic mtDNA variants is as high as 1:400 in diseases such as Mitochondrial encephalomyopathy, lactic acidosis, and stroke-like episodes (MELAS 3243A>G mtDNA mutation) (5, 6). In most disorders associated with variants in mtDNA, the individuals’ cells are heteroplasmic in that they contain a mixture of a variant and wild-type mitochondrial genomes (wt-mtDNA), with ratios that can also be tissue specific. The manifestation of defects associated with pathogenic mtDNA variants and the severity of symptoms depends on reaching of the proportion of variant mitochondria to certain threshold levels which is related to both nature of the variant and to tissues specific energy expenditure relying on mitochondria (7). Strategies to prevent transmission of mtDNA disorders include the use of donated oocytes, preimplantation genetic diagnosis, and mitochondrial replacement therapy (8-10), which involves the transfer of nuclear DNA from a heteroplasmic oocyte or embryo into a donor cytoplast with wild-type mitochondria by pronuclear or spindle transfer (9). Although successful in human, recent data have shown that incompatibility between donor and host mitochondria is associated with genetic drift leading to loss of donor mtDNA and reversion to the mutant haplotype (11, 12). An alternative approach to reduce heteroplasmy for variant mtDNA below threshold levels could be the use of mitochondria targeting nucleases that selectively cleave specific mtDNA haplotypes, resulting in their degradation and shifting the heteroplasmy ratio towards the wt mtDNA haplotype. Proof-of-principle for this concept has been demonstrated with restriction endonucleases (reviewed by (13)), mitochondrial zinc-finger nucleases (mtZFN) (14, 15) and mitochondrial transcription activator-like effector nucleases (mito-TALENs) (16-18). In contrast to restriction enzymes for which only a few applicable mutant sequences exist, mtZFN and mito-TALENs can be designed and engineered to selectively cleave a range of mtDNA sequences (19, 15, 20). Limitations to these approaches include the relatively labor and cost-intensive production, and achieving size compatibility with current virus-based delivery systems to tissues, and the need for repeated transfections to achieve effective heteroplasmy shift (17, 15). Another powerful genome editing and targeting methodology is based on clustered regularly interspaced short palindromic repeats (CRISPR) which is a part of bacterial immune systems that recognizes a 2-6 bp DNA sequence called protospacer adjacent motif (PAM) immediately following the DNA sequence targeted by nuclease (21). The most widely used constructs use the *CRISPR*-associated protein-9 nuclease (*Cas9*) from Streptococcus pyogenes (spCas9) and a chimeric single guide RNA (sgRNA) recognizing NGG as PAM sequence. Development of this and other emerging CRISPR based gRNA editing tools such as the Cpf1-family T-rich PAM sequence at the 5’ end as compared to the spCas9 recognition sequence 3’-NGG (22) would be very effective in editing mitochondrial DNA. However, a major obstacle in the development of CRISPR-Cas9 mediated gene editing of mitochondrial genome is the lack of effective delivery methods to deliver gRNA through mitochondrial membrane since the import of mRNAs into the mitochondria is largely dependent on tRNA (23-25). One approach to deliver nuclear encoded RNA into mitochondria involves the utilization of a native RNA transport enzyme, polynucleotide phosphorylase (PNPase) encoded by *PNPT1* gene, located in the mitochondrial inter-membranous space (26). There is increasing evidence of PNPase involved in the import of small RNAs, into mitochondria (27, 28). Further, it has been shown that addition of twenty nucleotide stem-loop sequence of nuclear RNAse P, to the transcripts not normally heading to mitochondria has been shown to facilitate transportation of these RNAs into mitochondria in a PNPase-dependent manner (29). However, this approach has not yet been demonstrated to deliver gRNAs into mitochondria.

Here, we hypothesized that PNPase-dependent RNA import can be used to transport sgRNAs into mitochondria and concurrently expression of Cas9 in the mitochondria through delivery of Cas9 mRNA with mitochondria localization signal would result in sequence specific cleavage of mtDNA in the cell reducing the heteroplasmy. We first designed sgRNA with stem loop sequence specific to A11204G region in the ND4 gene in the mtDNA and co-transfected them into the cells along with mitochondria targeting Cas9 constructs (MLS-Cas9). The results showed that the population of mtDNA carrying A11204G variant in their ND4 sequence was reduced remarkably and decreasing the ND4 expression in the cells. Further, fluorescent tagged gRNA sequence with RP-loop co-localized with mitochondria. This suggests that CRISPR/Cas9 targeting of mitochondria utilizing PNPase mediated transport is feasible and promise new opportunity for researchers to edit disease causing mutations in the mitochondrial genome.

## MATERIALS AND METHODS

### CRISPR/Cas9 Constructs

The expression construct pX330-U6-Chimeric_BB-CBh-hSpCas9 (Addgene; plasmid # 42230 (30)) for human codon-optimized SpCas9 and a chimeric guide RNA were modified as follows: The two nuclear localization signals (NLS) flanking the N and C terminals of Cas9 were replaced with two mitochondrial localization signals (MLS1 and MLS2); and (2) the human SpCas9 sequence was codon optimized for mouse expression (in the following referred to as mSpCas9). This modified expression plasmid is referred to as pX-U6-chimeric-MLS-mSpCas9. MLS1 consists of the amino-terminal leader peptide of mouse ornithine transcarbamylase (31)) and MLS 2 is of the 23-amino acid leader peptide of cytochrome oxidase subunit 8 (COX 8; (16)). The mSpCas9-MLS construct was synthesized by GenScript (NJ, USA) using their OptimumGene™ - Codon Optimization algorithm.

The region of mouse ND4 sequence with variant A11204G, that introduces a natural PAM sequence (AGG) for spCas9, was selected to design 19-20 bp target region of sgRNA using a CRISPR design online tool (http://crispr.mit.edu/guides), The selected sgRNA with additional 20 bp RP-loop [5’TCTCCCTGAGCTTCAGGGAG-3’] at the 5’ end of guide RNA was custom synthesized by Genscript, cloned into plasmid pUC57 with unique restriction sites (Pcil, Xba1), and then sub-cloned into pX-U6-chimeric-MLS-mSpCas9 to generate pX-U6-RP-sgRNA-MLS-Sp Cas9. (Fig. 1A). The same vector without the RP-loop sequence (pX-U6-sgRNA-MLS-mSpCas9) was used as control. Both constructs were then used as templates for in vitro RNA synthesis of sgRNAs with or without RP loop or Cas9 with MLS. RNA secondary structure predictions for RP Loop CRISPR were performed using *M-fold* software for recombinant sgRNA modeling. (Fig.1B, (32)).

**Figure 1.**
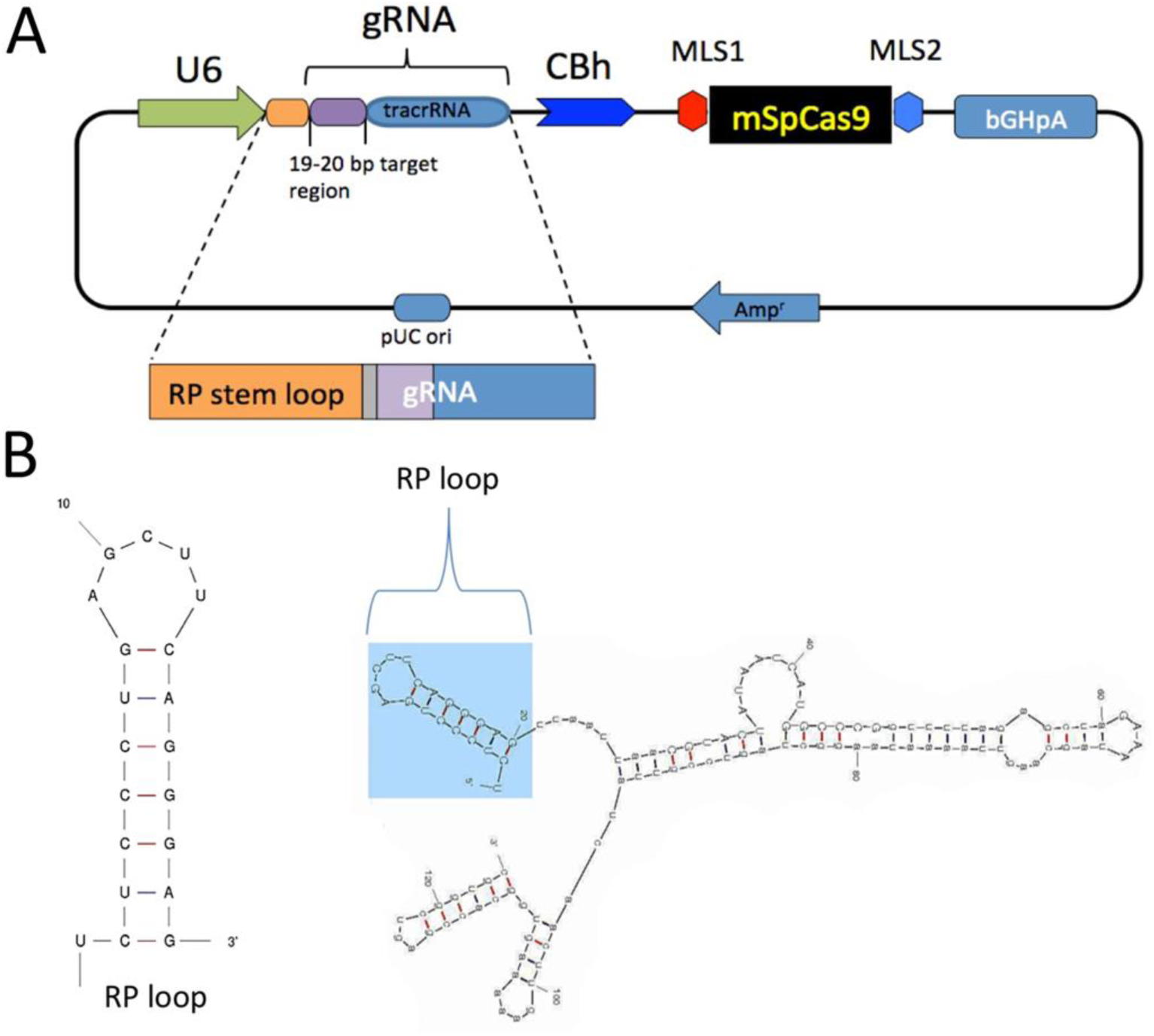
Generation and Structure of Mitochondria Targeting CRISPR-Cas9 Complex. Mitochondrial RP loop and gRNA targeting ND4 is in the downstream of U6 promoter and Chicken Beta Actin (CBh) controls the expression of mSpCas9 flanked by MLS1 and MLS2 sequences (**A**). M-Fold analysis of chimeric gRNA with RP loop (**B**).

### Generation of MLS-Cas9 mRNA and RP loop-sgRNA

The DNA template for in vitro transcription of mSpCas9 was generated by PCR amplification of pX-U6-RP-sgRNA-MLS-mSpCas9, using a forward primer that included a T7 promoter (T7 HAtag-Cas9-F: 5′TAATACGACTCACTATAGGGATGTACCCATACGATGTTCCAGATTACGCT-3′) and a reverse primer (Cas9-R: 5′-GCGAGCTCTAGGAATTCTTAC-3′). Cas9 mRNA (referred to as Cas9 in the following) was then synthesized, using the mMESSAGE mMACHINE® T7 Ultra Kit (Life Technologies, Carlsbad, CA) and purified by Lithium Chloride precipitation. In vitro RNA synthesis of sgRNA with and without RP-loop was also performed by using DNA templates of sgRNAs generated by PCR amplification of plasmid constructs, using construct specific forward primers that included a T7 promoter (pX-U6-RP-sgRNA-MLS-Sp Cas9, T7-RPloop-F: 5’TAATACGACTCACTATAGGGTCTCCCTGAGCTTCAGGGAGT-3’; pX-U6-sgRNA-MLS-Sp Cas9, T7-NoLoop-F: 5’TTAATACGACTCACTATAGGGCGTACTATAATCATGGCCCG-3’) and a common reverse primer (sgRNA-R: 5′-AAAAGCACCGACTCGGTGCC-3′). The RP loop sgRNAs were then synthesized using the MEGAshort-script™ T7 Kit (Life Technologies). RNA was purified and concentrated by using RNA Clean & Concentrator-5 Kit (Zymo Research Corp. Irvine, CA). The integrity of the synthesized RNAs was assessed using Agilent RNA 6000 Nano Kit with Agilent 2100 Bioanalyzer (Agilent Technologies, Santa Clara, CA), these newly synthesized sgRNA constructs will be referred to as RP Loop sgRNA (RP-loop sgRNA) and No RP Loop sgRNA (sgRNA).

### Cell culture

Primary mouse embryonic fibroblast (MEF) were derived from Tg(DR4) 1Jae/J mice stock No: 003208 (Jackson Laboratories). Human HEK293K cells were ATCC CRL-1573 (American Type Culture Collection, USA). Transient transfection with synthetic mRNA of Cas9 and sgRNAs was performed in either MEF or 293K cells using the TransIT®-mRNA Transfection Kit (Mirus Bio LLC) in OptiMEM medium (Invitrogen). For experiments with only wildtype ND4 MEFs being transfected with the CRISPR/Cas9 system (all except those in Fig. 3A) DMEM medium was supplemented with 50μg/ml of uridine and 100mM of pyruvate to improve cell survival after transfections.

**Figure 2.**
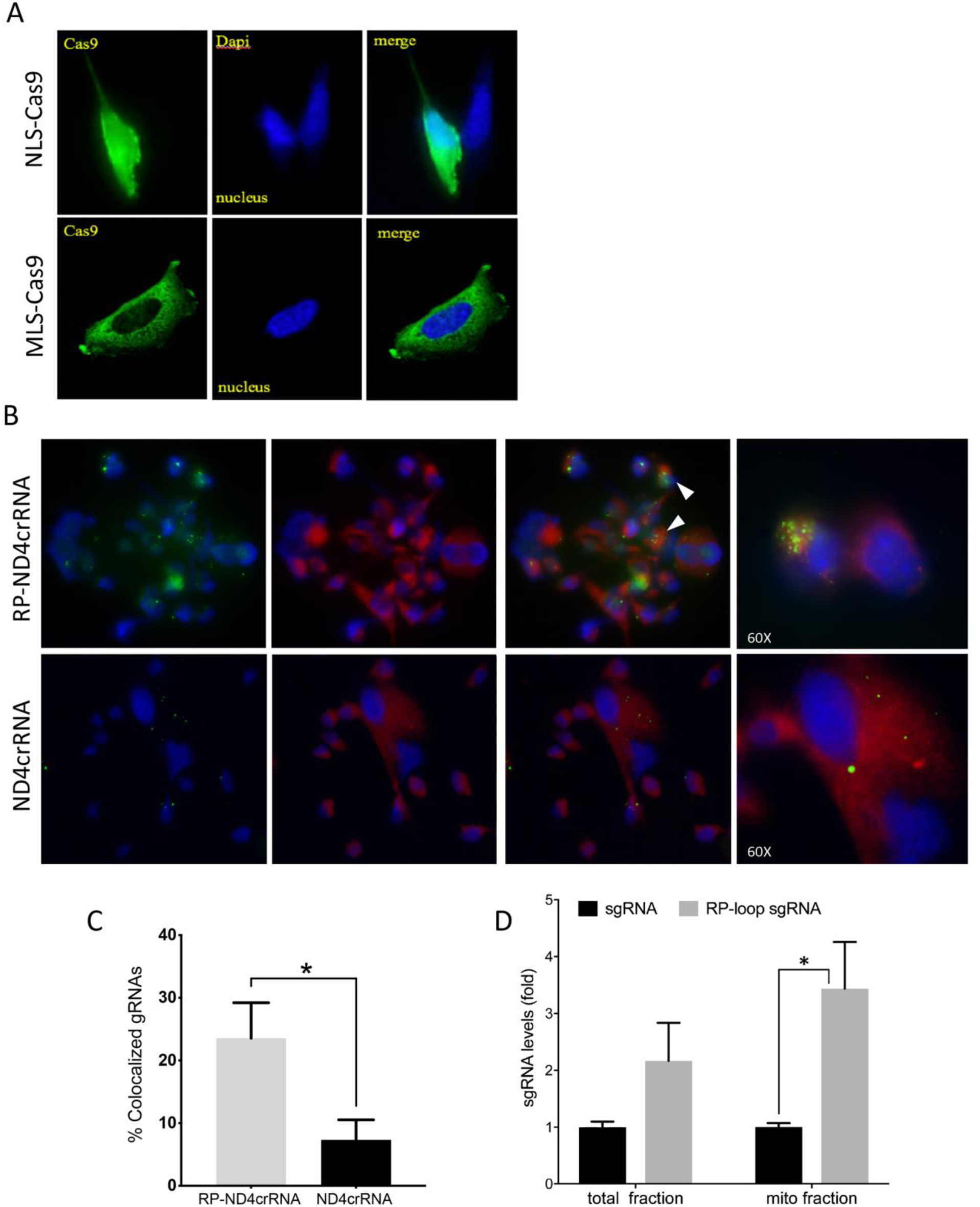
Delivery of CRISPR-Cas9 Complex into the mitochondria. (**A**) Immuno-staining showed at 24 hours post transfection MLS-Cas9 predominantly localized in the cytoplasm whereas NLS-Cas9 went to the nucleus (**B**) MEF Cells were transfected with Alexa flour 488 tagged ND4crRNA and RP-ND4crRNA and mitochondria stained after 18h with MitoTracker® Deep Red FM. Cells mounted in Vectashield anti-fade mounting media with DAPI (Vector Laboratories) and images taken at 20x and 60x magnifications using an Olympus BX41 microscope and SPOT camera (Olympus, Japan) to observe co-localization. (**C**) Cells with mitochondrial co-localized guideRNA (solid arrows) were counted at 20x magnification showed enhanced percentage of RP-ND4crRNA merged with mitochondria (n=3, Student’s t-test p=0.01). (**D**) RP-loop increases the levels of sgRNA detected in the mito fractions. qPCR was performed and levels of sgRNA was normalized to mitochondrial ND4 and ND1 as internal controls and reported as fold change 2^ΔΔCT^(n=3, Student’s t-test, p<0.05).

**Figure 3.**
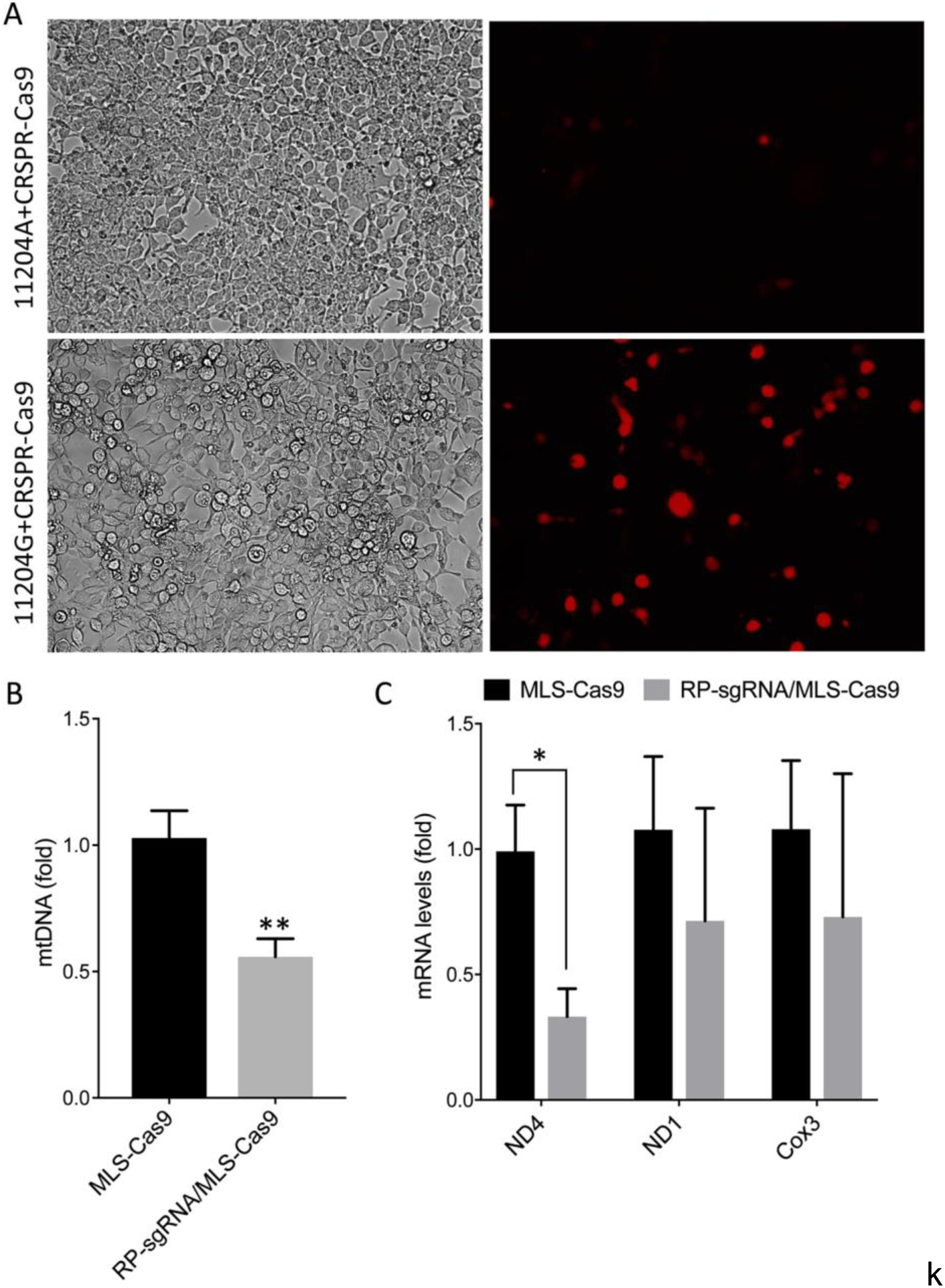
Specificity and functionality of RP-loop CRISPR/Cas9 complex. (**A**) Specificity of RP loop-CRSPR-/Cas9 was determined using the GeneArt^®^ Genomic Cleavage Selection Kit that expressed orange fluorescent reporter, in 293K cells 72 h post transfection, specific to 11204 G variant of ND4. qPCR for mtDNA levels (**B**) qRT-PCR for ND4 mRNA expression **(C)** was significantly lower in RP-loop sgRNA-CRISPR/Cas9 transfected MEF cells, at 24 hours post transfections, compared to MLS-Cas9 controls. (n=5, Student’s t-test *p=0.016; **p=0.005).

### *In vitro* Assay to Test mito-CRISPR Functionality and Specificity

Mito CRISPR target selection was performed using the GeneArt® Genomic Cleavage Selection Kit (cat# A27663 ThermoFisher Scientific), which is based on restoration of reporter Orange Fluorescent Protein (OFP) expression if endonuclease activity at a target sequence induces DNA double-stranded break and repair. Target DNA fragments containing the sgRNA target sequence with one nucleotide difference (112004G) and control (112004A) in ND4 fragment were cloned separately into the pGCS reporter vector producing pGCS-wt and pGCS-variant. Co-transfection of reporter constructs into HEK293K was performed using TransIT-2020 transfection reagent (Mirus Bio LLC), with RP-sgRNA/Cas9. Orange fluorescence was visually assessed at 48 to 72 hrs. post transfection using an EVOS cell imaging system (ThermoFisher Scientific, USA).

### Immunostaining

MEF cells were grown on cover slips in 12-well plates and transfected with either pX330-U6-Chimeric_BB-CBh-hSpCas9 or pX-U6-chimeric-MLS-mSpCas9. After 24 h, medium was removed and cells fixed by 2 brief washes in ice-cold acetone. Cells were blocked in 3% goat serum in 1xTBS for 30 min, followed by immunostaining with Cas9 monoclonal antibody 4G10 (Diagenode cat#: C15200216; 1:200 dilution) with gentle shaking for 2h at room temperature. Cells were washed in 1% goat serum in 1xTBS, followed by incubation with secondary anti-mouse antibody conjugated with Alexa 488 (Molecular Probes) 1:1000 dilution for 1h at room temperature. Cells were washed with 1.0% goat serum in 1xTBS cells and mounted in Vectashield with DAPI and images taken using confocal microscope (Zeiss LSM 700).

### Mitochondrial DNA Isolation

Mitochondrial DNA was isolated using Mitochondrial/Cytosolic Fractionation Kit (BioVision Inc., CA; cat# K256-25) according to the manufacturer’s instructions. MEF cells were grown in 6-well plates to 80% confluency before transfection with RP-sgRNA11204G (1.5ug/well) using TransIT®-mRNA Transfection Kit (Mirus Bio LLC). After 24h post transfection, 5×10^6^ cells were harvested using trypsin (0.05% trypsin-EDTA). Cell membranes were disrupted in cytosolic buffer using a dounce homogenizer followed by successive centrifugation steps at 700xg to collect supernatant followed by 10,000 x g centrifugation to collect intact mitochondria. Mitochondrial pellets were then used for isolating mito-DNA using QIAamp DNA mini Kit (Qiagen). DNA was eluted in water and quantified by NanoDrop 2000 UV-Vis Spectrophotometer (Thermo Scientific).

### Quantitative PCR

DNA and RNA were isolated using QIAamp DNA Mini Kit (Qiagen) and mirVana RNA isolation kit (Ambion-ThermoFisher), respectively. cDNA synthesis of RNA was performed by using SuperScript® VILO™ cDNA Synthesis Kit (ThermoFisher Scientific; cat# 11754050). QPCR was performed using Precision Melt Supermix containing EvaGreen dye (cat# 172-5110) using CFX96 Touch™ Real-Time PCR Detection System (BioRad, USA). Sequences of PCR primers are listed in Table.

**Table:**
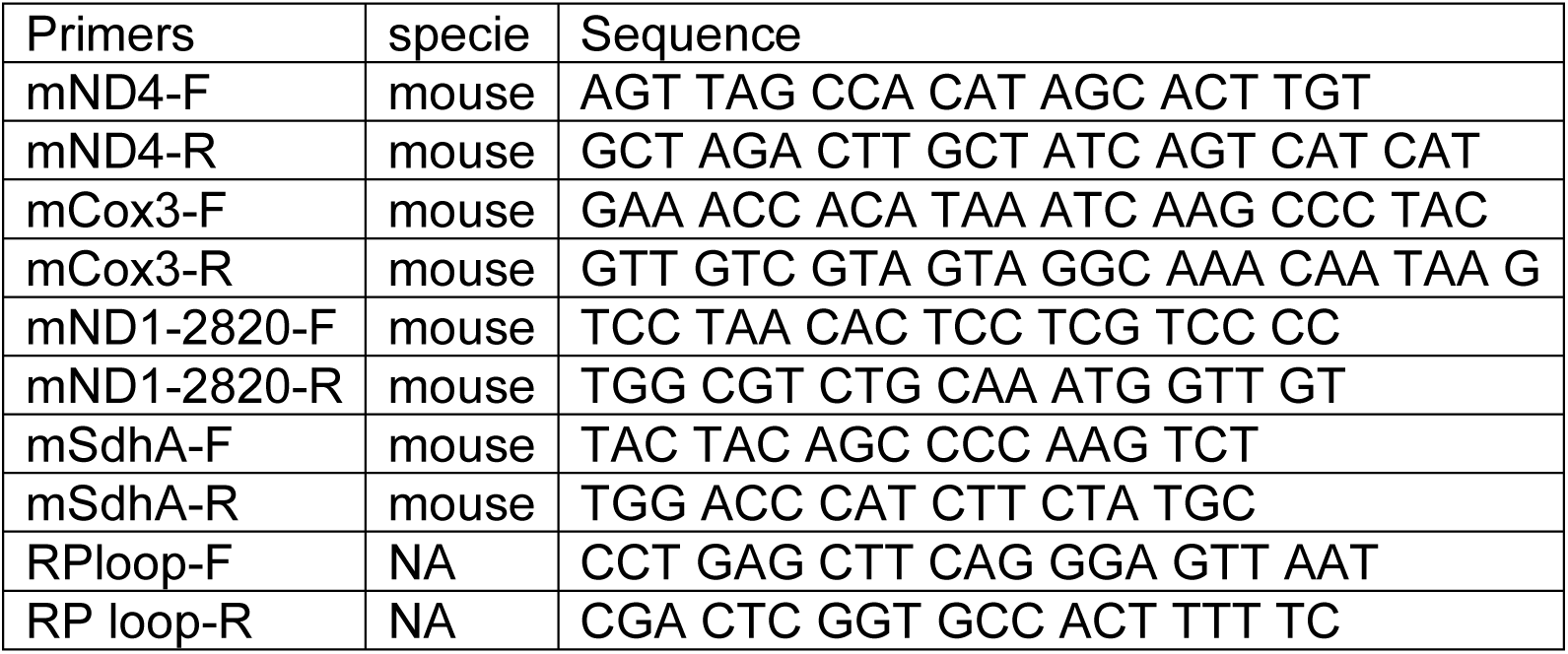
Primers for qPCR and RT-qPCR.

## RESULTS

### mSpCas9 Do Not Localize to Nucleus

To facilitate transport of Cas9 into the mitochondria of mouse cells, we modified a CRISPR/Cas9 expression plasmid (pX330-U6-Chimeric_BB-CBh-hSpCas9 (30)) such that a mouse-optimized Cas9 sequence was flanked by two mitochondrial localization signals (MLS) (Fig. 1A). The amino terminal signal (MLS1) consisted of the amino-terminal leader peptide of mouse ornithine transcarbamylase (31) and the C-terminal MLS2 consisted of the 23 amino acids leader peptide of cytochrome oxidase subunit 8 (16). Because codon bias can affect translation and activity of Cas9 protein in cell culture systems (33) we also used a Cas9 coding sequence that was optimized for mouse expression. As expected, transfection of MEF cells with pX330-U6-Chimeric_BB-CBh-hSpCas9 (30)), in which the coding sequence of Cas9 is flanked by nuclear localization signals, resulted in strong nuclear signal of Cas9 in immunostaining assays (Fig. 2A top panel). In contrast, cells transfected with the modified expression construct encoding Cas9 flanked by MLS1 and MLS2, lacked Cas9 signal in the nuclei and exhibited immunostaining throughout the cytoplasm. We did not perform mitochondrial staining for co-localization as the two leader peptides used in our construct are well known for delivery into mitochondria and the objective of performing Cas9 IHC was only to confirm that by replacing NLS with MLS we have indeed prevented nuclear translocation of this modified Cas9 suggesting mitochondrial localization (Fig. 2A bottom).

### Guide RNA Delivery into Mitochondria

To determine whether RP loop mediates import of guideRNA construct into mitochondria, we used a 20-nucleotide stem-loop element (RP-loop) that is a component of nuclear RNAse P at the 5’end of guide. PNPase is localized to the inner mitochondrial membrane and regulates the import of nuclear-encoded RNAs into the mitochondrial matrix. Addition of the RP-loop to transcripts that do not normally translocate to mitochondria has been shown to allow for RNA import in a PNPASE-dependent manner (29). Furthermore, this delivery system has been previously reported to mediate the targeted transfer of recombinant RNAs into mitochondria (29, 34). We therefore constructed a hybrid sgRNA in which the RP loop was appended to the 5’-end of the sgRNA11204G construct designed to selectively base-pair with the wild-type mtND4 target -20 nucleotide sequence. To facilitate future sub-cloning, an 8-nucleotide Pac1 restriction site separated the RP-loop and sgRNA (Fig. 1A). Based on structure predictions using the M-fold algorithm, the hybrid RP-loop-gRNA maintained the secondary structure of the stem-loop required for mitochondrial import [Initial ΔG= -42.60 kcal/mole at 37 C] (Fig. 1B).

To test if the RP-loop sgRNA can localize to mitochondria we labeled both, RP-loop sgRNA and sgRNA without RP-loop, with Alexa flour 488 at 3’ end. However, due to size limitation for the synthesis of fluorescent labeled RNA we truncated the tracrRNA sequence and named these truncated guideRNA as RP-ND4crRNA (Sequence: 5’rUrCrUrCrCrCrUrGrArGrCrUrUrCrArGrGrGrArGrUrUrArArUrUrArArCrGrUrArCrUrArUrArArU rCrArUrGrGrCrCrCrGrGrUrUrUrUrArGrArGrCrUrAr GrArArA/3AlexF488N/) and ND4crRNA (5’rUrUrArArUrUrArArCrGrUrArCrUrArUrArArUrCrArUrGrGrCrCrCrGrGrUrUrUrUrArGrArGrCr UrArGrArArA/3AlexF488N/). We transfected RP-ND4crRNA and ND4crRNA into MEF cells and after 18hrs stained mitochondria in live cells with MitoTacker Deep Red FM (Cat# M22426, Invitrogen). We observed a significantly higher number of cells with RP-ND4crRNA that co-localized with mitochondria relative to ND4crRNA (p<0.01) (Fig. 2B & C)

Further, as an added evidence of mitochondrial localization of the RP-loop sgRNA construct, we transfected RP-loop sgRNA11204G and sgRNA11204G into MEF cells and isolated mitochondrial fractions after 24h. qPCR showed that though both full length RP-loop guide RNA and sgRNA were present in mitochondrial fraction, RP-loop sgRNA was highly enriched in mitochondrial fraction of transfected cells versus sgRNA controls (Fig. 2D) indicating that RP loop added sgRNAs are efficiently transported into the mitochondria. The presence of sgRNA without RP loop in mitochondrial fraction may be due to unregulated transport between inner and outer mitochondrial membrane that has earlier been proposed to facilitate CRISPR/Cas9 based editing in mitochondria (35).

### Targeting Mitochondrial DNA using CRISPR/Cas9

To evaluate CRISPR/Cas9-mediated cleavage of a mitochondrial-encoded gene, we used mouse haplotypes of the mitochondrial-encoded NADH:ubiquinone oxidoreductase core subunit 4 (mtND4) that had been generated in mouse cell lines (36). Specifically, we targeted residue 11204 of mtND4, at which a G->A variant of residue 356 (R356Q) corresponds to a mutation in human MT-ND4 associated with complex I deficiency and respiration defects (36). Effectively this approach would be useful to enrich for a variant in a heteroplasmic cell by eliminating or reducing the fraction corresponding to the disease causing variant. Using mouse mtDNA sequences, we selected target guide sequences against base 11204G of mt-ND4, which encodes R356, and used these to construct a hybrid guide RNA (sgRNA11204G) composed of the CRISPR array and tracrRNAs (30). The guide sequence selected (5’CGTACTATAATCATGGCCCG-3’) scored 92 on the scale of zero to 100 to indicate the faithful on-target activity of guide with only 1 off-target site. We assessed activity and allele-specificity of sgRNA11204G in vitro in HEK293K cells following co-transfection with Cas9 expression plasmid and orange fluorescent protein (OFP)-based reporter constructs for mtND4 112004G and mtND4 112004A. OFP fluorescence, indicating base-specific cleavage, was observed in cells transfected with the 11204G sequence but not with the mutant 11204A sequence (Fig. 3A).

### Inclusion of the Stem Loop Facilitates CRISPR-Cas9 Mediated Reduction of mtDNA with A11204G variant

With the inclusion of MLS sequences, Cas9 can be translocated to the mitochondria. However, functional localization within the mitochondria requires evidence of targeted mtDNA endonuclease activity. To address this, we co-transfected wild-type MEFs with a MLS-Cas9 expression construct either as mRNA or as plasmid alone or in combination with RP-sgRNA11204G. The CRISPR/Cas9 complex with RP-loop sgRNA11204G was able to significantly reduce mtDNA levels relative to Cas9 alone (P=0.005; Fig. 3B). This knockdown of mtDNA levels post 24h by MLS-Cas9 and RP-sgRNA11204G was significant when compared to the MLS-Cas9 only treated group. To investigate if mtDNA knockdown downregulate ND4 gene expression, we next quantified expression levels of mitochondrial-encoded transcripts in only wild-type MEF that has 11204G in ND4 sequence. At 24h post-transfection, we observed a significant reduction of ND4 transcripts in the RP-sgRNA11204G group when compared to transfection with the sgRNA11204G group, using Gapdh as internal nuclear RNA control (Table, Fig. 3C). Other mitochondrial genes such as ND1 and Cox3 also had lower transcripts levels but were not significant when compared to the sgRNA11204G treatment group, this lower level may reflect the lower levels of mtDNA overall due to CRISPR/Cas9 mediated cleavage of mitochondrial genome (Fig. 3C) that has been shown to rapidly degrade in vivo (37, 38).

### Inclusion of stem loop is critical for knock-down of mitochondrial genes

To evaluate if the RP-sgRNA11204G increased mitochondrial translocation leads to an increase in the target gene knockdown we mixed (1:1) fibroblast cell line carrying either 11204G ND4 target and a variant fibroblast cell line with 11204A ND4 sequence, a gift from Dr. M F Alexeyev at University of South Alabama, USA. Ideally a cybrid cell line would be the best approach to determine in vivo specificity of the sgRNA as an alternative we used this pseudo-heteroplasmic system to evaluate the specificity of CRIPSR/Cas9 on the target mitochondrial sequences. The cells were transfected with CRISPR/Cas9 complex with or without RP-loop sgRNA. Twenty-four hours post transfection, we found that only RP-sgRNA that is specific for the variant 11204G caused a significant reduction in the ND4 with 11204G sequence when compared to control group, sgRNA11204G with Cas9 was not capable of reducing the ND4 levels indicating that RP loop is essential for mitochondrial gene silencing. (Figure 4).

**Figure 4.**
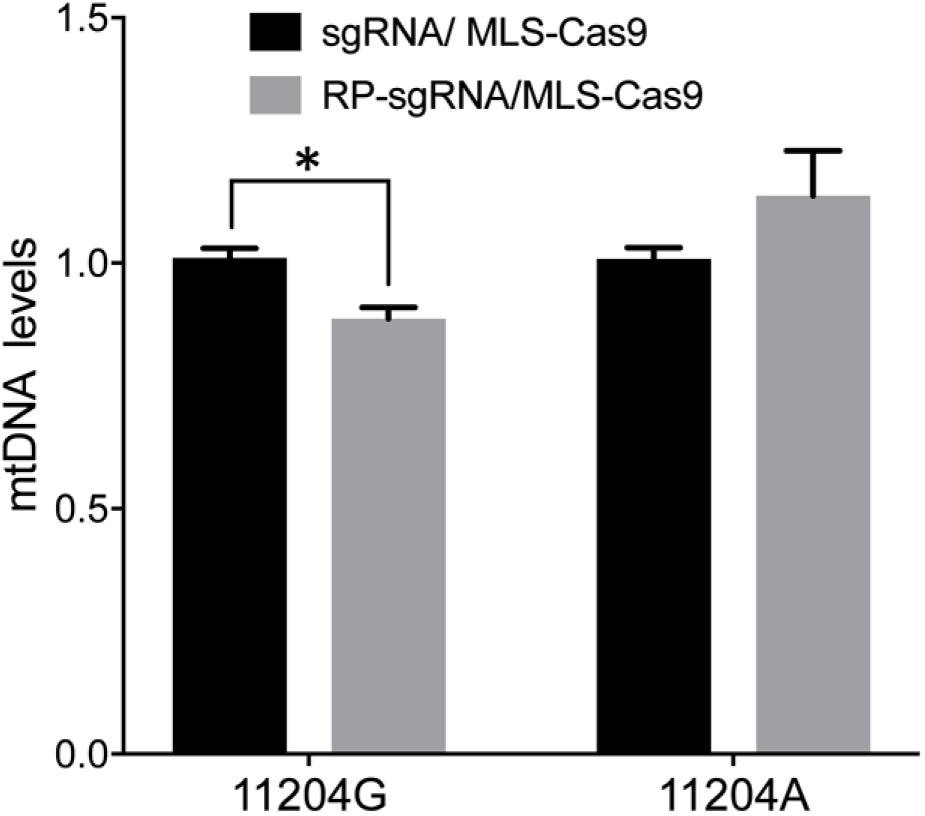
RFLP analysis of ND4 variants targeted by CRISPR-Cas9 complexes. Specific reduction of ND4 11204G variant by RP-loop sgRNA/MLS Cas9 complex in sequence specific manner after transfection into with 50:50 mixture of fibroblast carrying either 11204G or 11204A ND4 variants. Restriction Fragment Length Polymorphism (RFLP) analysis was performed by PCR amplification of 941bp ND4 fragment (RFLP-ND4-F 5’atttgaagcaaccttaatccca; RFLP-ND4-R 5’ttagttctcgtgtgtgtgaggg) with a unique Ava1 site present in 11204G variant. Digested DNA was resolved on 1.2% agarose gel stained with ethidium bromide. Band intensity for the uncut (941bp) and the Ava1 cut fragment (627bp and 269bp) was quantified by ImageJ 1.50g image processing program. A reduction in band intensity, specifically of cut fragment with the RP-loop appended sgRNA only was observed (n=3, Student’s t-test p<0.05).

## DISCUSSION

Our objective was to determine whether it was possible to enable or enhance CRISPR/Cas9 mitochondrial genome editing utilizing PNPASE a native RNA transport system although the mechanism of augmented import of RNA by PNPASE is not fully understood (39). We conclude that both the targeted mtDNA and its transcripts levels are reduced and correlate with the designed mitochondrial import of the target specific guide RNA. Previous reports of CRISPR based editing (22) have shown similar levels of reduction in targeted mtDNA and transcripts without modifying the gRNA, however we only observe this in our modified sgRNA specific for the target sequence allowing uptake into the mitochondria via PNPase. The differences may be attributed unregulated uptake of small RNA into mitochondria or the difference in transfection method, incubation time and repeated transfections. One of the critical experiment for the proof-of-concept was to visually show that gRNA can be imported into mitochondria if RP-loop is appended. Therefore, we selected Alex flour 488 to label the construct. The advantage of using Alex Fluor 488 is that unlike other fluorescent dyes like Cy3, this dye does not accumulate at mitochondrial surface (40) therefore, we posit that if RP-loop is guiding RNA then visualization of Aexa488 labeled gRNA to mitochondria would be possible. Our results show an enhanced delivery of gRNA with RP-loop. A recent study has shown that guide RNA with short hairpin structures can promote mitochondrial import and specific cleavage of mtDNA albeit at low level due to limited import into mitochondrial matrix (41, 42). This study further strengthens our hypothesis that specific hairpin structures that are involved in the delivery of nuclear encoded RNA can serve as potential adapter for delivery and PNPase may serve as conduit for channeling this recombinant RNA into mitochondrial matrix. The presence of multiple mitochondrial genomes in each cell may result in only a cleavage of a subset of mitochondrial genomes with variable impact on viability of each mitochondria. Presumably repeated treatment or longer exposure times would correspond with an increase in mitophagy in competition with replication of non-targeted mitochondrial genomes. Further, a rapid reduction in mtDNA copy number as shown with previous studies using mito-TALEN creates potential risk of developing mtDNA depletion syndrome which limits the clinical implication (43).

For all our studies with knockdown, the DNA or RNA was isolated 24h after transfection, at a point that we found an optimal choice for loss of cell viability and mitochondrial gene expression. To avoid cell death, we used in vitro synthesized RNAs for both sgRNA and Cas9 instead of using plasmid which would continuously express uncontrolled levels of the complex. Moreover, this approach only reduces ∼25% of mtDNA which would avoid any mitochondrial depletion syndrome. Our present study provides support for the function of the RP loop as an enhanced mitochondrial transporter of gRNA for CRISPR-Cas9 machinery across the mitochondrial membranes.

Currently validation of alternative endonuclease based systems including TALENS and ZFN is ongoing in clinical trials and presently have the advantage of a large range of targetable nucleotides within the mitochondrial genome. However, the main appeal for using CRISPR/Cas9 endonuclease is ease of design and speed of execution. With the introduction of other variants of Cas9 such as saCAs9, Cpf1 which performs cutting functions analogous to spCas9 but have different PAM recognition sequence and smaller size of endonuclease, there will be more efficient delivery to the cells and mitochondria. We anticipate that our proof-of-concept study with chimeric guideRNA with RP-loop may be also able to deliver other CRISPRs to the mitochondria this will increase the available target sequences in the mitochondrial genome for potential therapeutic option by shifting heteroplasmy.

We observed relatively robust RP loop mediated mitochondrial localization of sgRNA in the mitochondria potentially mediated via PNPase present in inner mitochondrial membrane and with MLS-Cas9 in the mitochondria the CRISPR/Cas9 complex promoted mtDNA knockdown (Figure 5). We can state that our report warrants for further investigation of this approach. Regardless, the utility of the RP loop mediated RNA transport versus other modes is only relevant if the engineered heteroplasmic shift is sufficient to address the symptomatic manifestation of mitochondrial diseases.

**Figure 5.**
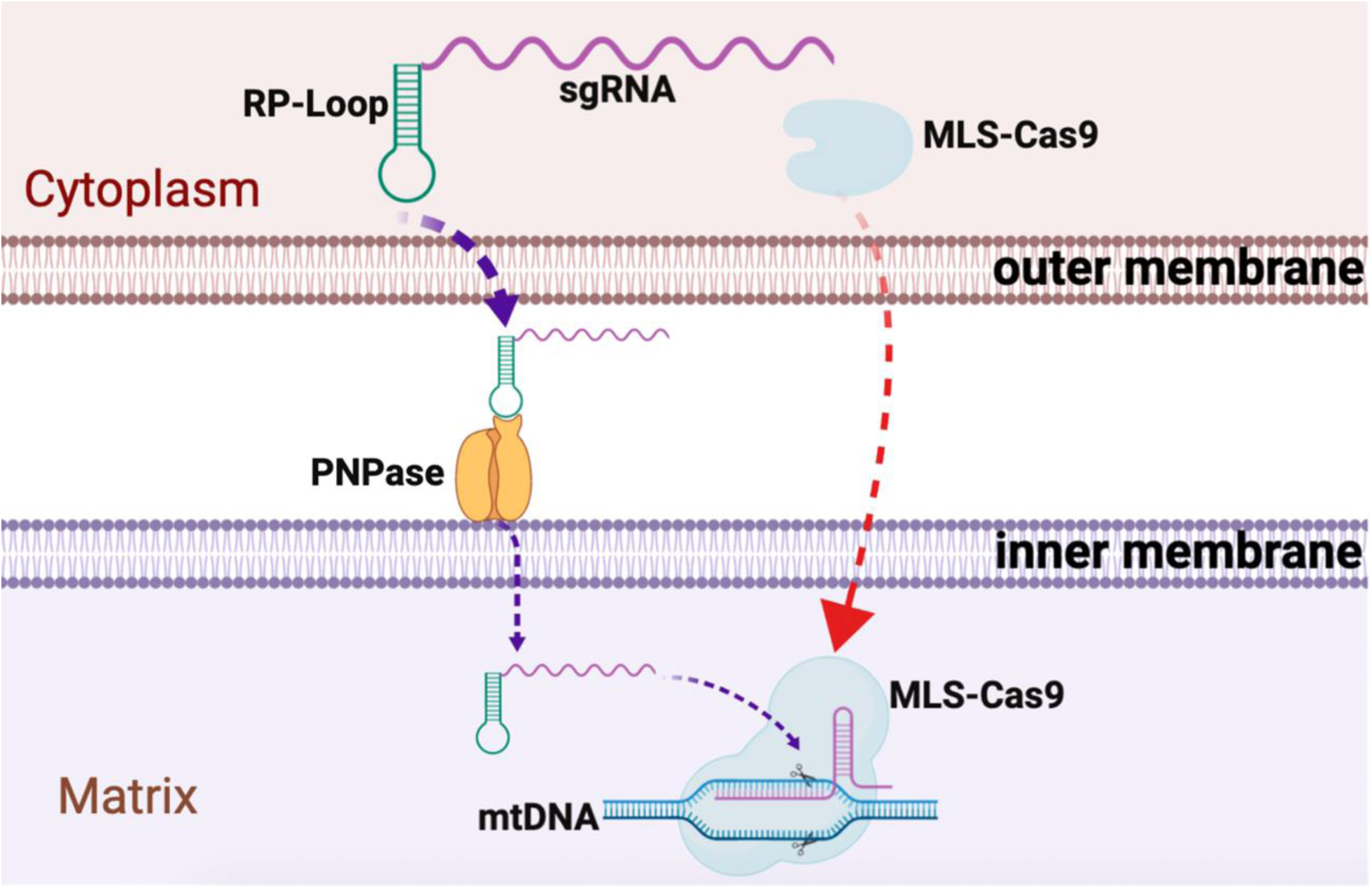
Graphical summary of CRISPR/Cas9 delivery into mitochondria. Guide RNA, with RP-loop attached, upon transfection is imported into mitochondria by attaching to PNPase in inner mitochondrial membrane (IM). Cas9 protein with N-terminus mitochondria localization sequences (MLS) targets Cas9 to mitochondria where it forms complex with RP-loop sgRNA to cause sequence specific mtDNA cleavage.

## AUTHOR CONTRIBUTIONS

SRAH and KJM conceptualize the project, designed research and wrote manuscript. SRAH and BK performed experiments. MEY and SE helped in writing and discussion. All authors approved the manuscript.

## ACKNOWLEDGMENT

The financial support for this study was through Nationwide Children’s Hospital Technology Development Funds to SRAH and KJM.

